# Disrupting the interaction between AMBRA1 and DLC1 is a promising therapeutic strategy for neurodegeneration that prevents apoptosis while enhancing autophagy and mitophagy

**DOI:** 10.1101/2023.10.13.562240

**Authors:** Kate Hawkins, Meg Watt, Sébastien Gillotin, Maya Hanspal, Martin Helley, Jill Richardson, Nicola Corbett, Janet Brownlees

## Abstract

Activating molecule in Beclin1-regulated autophagy (AMBRA1) has critical roles in autophagy, mitophagy, cell cycle regulation, neurogenesis and apoptosis. Dysregulation of these processes are hallmarks of various neurodegenerative diseases and therefore AMBRA1 represents a potential therapeutic target. The flexibility of its intrinsically disordered regions allows AMBRA1 to undergo conformational changes and thus perform its function as an adaptor protein for various different complexes. Understanding the relevance of these multiple protein-protein interactions will allow us to gain information about which to target pharmacologically. To compare potential AMBRA1 activation strategies we have designed and validated several mutant constructs (ACTA, TAT, WD40 and S1014) in addition to characterising their effects on proliferation, apoptosis, autophagy and mitophagy in SHSY5Y cells. AMBRA1^TAT^, which is a mutant form of AMBRA1 that can’t interact with dynein light chain (DLC)1 at the microtubules, produced the most promising results. Indeed, overexpression of this mutant protected cells against apoptosis and induced autophagy and mitophagy in SHSY5Y cells in addition to enhancing the switch from quiescence to proliferation in mouse neural stem cells. Future studies should focus on designing compounds that inhibit the protein-protein interaction between AMBRA1 and DLC1 and thus have potential to be used as a drug strategy to treat neurodegeneration.

## Introduction

AMBRA1 is an intrinsically disordered protein (IDP) with a molecular mass of approximately 130kDa. IDPs participate in many biological processes, and the flexibility of their disordered regions allows for conformational changes and thus binding to various different proteins depending on the cellular context. AMBRA1 acts as a scaffolding protein in this way to form complexes to facilitate proliferation, apoptosis, autophagy and mitophagy (Cianfanelli et al., 2015, Di Leo et al., 2021, Di Rita et al., 2018b, Chaikovsky et al., 2021). Interestingly, these processes (particularly autophagy) have been implicated in neurodevelopment and adult neurogenesis (Gillotin et al., 2021). Mutations in AMBRA1 cause neural tube defects in both mice (Fimia et al., 2007) and humans (Ye et al., 2020) and downregulation of AMBRA1 has been shown to lead to reduced survival of neural stem cells (NSCs) (Yazdankhah et al., 2014, Vazquez et al., 2012). In addition, one group observed enhanced reactivation of quiescent NSCs in BECLIN1/BCL2 mutant mouse NSCs in which BECLIN1 is constitutively active (Wang et al., 2022). However, this is yet to be shown for AMBRA1 specifically.

The main role of AMBRA1 is in the autophagy signalling network, which mediates lysosomal degradation of damaged or redundant cellular components (Aman et al., 2021). When autophagy is initiated, for example by nutrient depletion, mTOR is inactivated thus allowing ULK1 activation. ULK1 then phosphorylates BECLIN1 and AMBRA1 to promote AMBRA1 release from the microtubules where it is tethered via dynein light chain (DLC)1, and subsequent activation of the VPS34 complex. The BECLIN1/VPS34 complex is then directed to the ER where it participates in autophagosome formation from the ER membrane (Fimia et al., 2011, Tang et al., 2016). In addition, a positive feedback mechanism occurs whereby AMBRA1 stabilises ULK1 by assisting TRAF6 to carry out K63 chain ubiquitination of ULK1, resulting in its self-association. Upon autophagy termination, other E3 ligases such as CULLIN4 and RNF2 (Xia et al., 2014) interact with AMBRA1 to mediate its degradation. For instance, CULLIN4 in association with DDB1 binds to AMBRA1 via its WD40 domain (Antonioli et al., 2014).

AMBRA1 also plays a key role in both PARKIN dependent and independent mitophagy. In the former process, AMBRA1 associates with the E3 ligase PARKIN to mediate ubiquitination and subsequent degradation of outer mitochondrial membrane proteins and in the latter the HUWE1 E3 ligase performs this function in the place of PARKIN. It has been shown that AMBRA1 directly binds to LC3 and that, during PARKIN independent mitophagy, IKKα induces phosphorylation of S1014 which stabilises the interaction between AMBRA1 and LC3 (Di Rita et al., 2018b).

Currently, there is no drug that has been described to activate AMBRA1. However, tagging a mitochondrial localisation sequence onto the AMBRA1 protein (AMBRA1^ACTA^) has been shown not only to induce mitophagy but to counteract the oxidative stress and apoptosis induced by either rotenone or 6-OHDA in SHY5Y cells (Di Rita et al., 2018a). This represents an interesting mechanism that could mimic some therapeutic strategies currently developed for neurodegeneration.

We have designed and overexpressed in SHSY5Y cells (which otherwise express negligible levels of AMBRA1 protein) various mutant AMBRA1 constructs representing potential activation strategies (Table S1). Following validation of these constructs we have characterised their effects on proliferation, apoptosis, autophagy and mitophagy. AMBRA1^TAT^ (which cannot bind to DLC1 at the microtubules) was shown to be the most promising mutant, inducing a clear switch away from apoptosis and towards autophagy and mitophagy. We then went on to show that this mutant construct could enhance the transition from quiescence to proliferation in mouse neural stem cells (mNSCs) as a model of neurogenesis. Together, these findings suggest that devising pharmacological strategies to target the AMBRA1/DLC1 interaction may be a promising strategy to tackle known impaired signalling pathways in neurodegeneration.

## Materials and Methods

### Cell culture

HEK293T (Sigma, 1202200) and SHSY5Y cells (Sigma, 94030304-1VL) were purchased from ECACC. SHSY5Y cells were cultured in Dulbecco’s modified Eagle’s medium (DMEM, Gibco, 31966-021) whereas HEK293T cells were cultured in DMEM (Gibco, 11330032). Both were supplemented with 10% FBS (Gibco, 11580516) and 1% penicillin-streptomycin solution (P/S, Gibco, 11548876). Cells were cultured at 37°C with 5% CO2.

Mouse neural stem cells (mNSCs) were purchased from R&D systems (NSC002). The cells were grown in proliferative conditions in complete basal media: DMEM/F12 + HEPES + Glutamine (Gibco, 11330032), 0.5X B27 supplement (Gibco, 1530536), 0.5X N2 supplement (Gibco, 17502048) 0.1 mM 2-Mercaptoethanol (Gibco, 3150010), 1X MEM NEAA (Gibco, 11140050), 7.5% BSA (Thermo, 15260037), 1.45 g/L D-(+)-Glucose (Sigma, G8644) and 1X P/S. The basal media was supplemented with 10 ng/mL EGF (Peprotech, 315-09), 10 ng/mL FGF (Peprotech, 100-188) and 1 mg/mL laminin (Sigma, L2020) to grow cells in proliferative conditions. To induce quiescence, plated cells were transferred into complete basal media supplemented with 0.05 ng/μL BMP4 (R&D systems, 5020-BP), 20 ng/mL FGF and 1 mg/mL laminin (Martynoga et al., 2013).

### Lentivirus generation and transduction

Mutant AMBRA1 lentiviral plasmids (Table S1) were generated by GeneArt Gene Synthesis (Thermo). HEK293T cells were transfected with lentiviral plasmids and the appropriate packaging plasmids using turbofectamine (Origene, TR30037) according to the manufacturer’s instructions. After 24 hours, the medium was changed to normal HEK293T media. After an additional 24 hours the media was either filtered onto SHSY5Y receiver cells or 10ml was added to 2.5ml PEG-IT reagent (Abcam, ab1025238) and stored at 4°C overnight. This viral supernatant was then concentrated by centrifugation at 3200 x g for 30 minutes at 4°C before being resuspended in 100 µl virus suspension solution (Abcam, ab102538), aliquoted and stored at -70°C. For transduction, 20 µl of this concentrated virus was added to 660 µl normal media which was added to a well of a 6 well plate of cells and left overnight before topping up the media with 2 mL the next morning.

### Western blot analysis

For autophagy flux experiments, cells were treated with 100 nM bafilomycin (Sigma, B1793) in normal growth media for 4 hours at 37°C. Cells were washed with PBS (Gibco, 15326239), detached with TrypLE and lysed in RIPA buffer (Thermo, 10017003) supplemented with protease inhibitors (Sigma, 4693116001) on ice for 15 minutes with intermittent vortexing. Samples were then centrifuged at 14,000rpm for 5 mins at 4°C before the pellet was discarded. Protein concentrations were determined with a Pierce™ BCA assay protein kit (ThermoFisher, 23225). The required sample volume was made up to 100 μL using sample, dH_2_O with NuPAGE LDS sample buffer (4X; Invitrogen, NP0007) and NuPAGE sample reducing agent (10X; Invitrogen, NP0004). Cell extracts were separated by sodium dodecyl sulphate–polyacrylamide gel electrophoresis (Novex gels cat. NP0335, SDS–PAGE). The membranes were blocked in PBS blocking buffer (LI-COR, 927-70001) for 30 minutes at room temperature before being immunoblotted using primary antibodies at their optimal dilution (Table S2, LC3-II for autophagy flux experiments) in PBS blocking buffer for approximately 4 hours at room temperature and then overnight at 4°C. The membranes were then washed with PBS and 0.1% Tween-20 (Sigma, P1379-100) before being incubated with secondary antibodies (Table S2) for 1 hour at room temperature and washed again. Immunoreactive bands were imaged using Odyssey CLx (LI-COR Biosciences). The band intensity was semi-quantified by normalising with an endogenous control protein (GAPDH) using ImageStudio Lite (Version 5.2.5 9; LI-COR Biosciences).

Autophagy flux was measured by quantifying the dynamic change in LC3-II expression following bafilomycin treatment. Bafilomycin causes a late-stage blockage in autophagy, inhibiting autophagosome lysosome fusion, which results in LC3-II accumulation. LC3-II expression, normalised to the endogenous control, was quantified and the difference between the bafilomycin treated and untreated samples created an autophagy flux score. This score was then used to compare basal autophagy flux between cells expressing various AMBRA1 mutants.

### Co-immunoprecipitation (CoIP)

SHSY5Y cells were pelleted and solublised at 4°C for 30 mins. Insoluble material was then removed by centrifugation at 1200rpm for 5 mins. AMBRA1 antibody (Proteintech, 1376-1-AP) was then coupled to dynabeads according to the manufacturer’s instructions (Thermo, 14311) before protein lysate was mixed with the beads. This was followed by multiple wash steps to collect the purified protein complex. The protein concentrations of the samples were then determined using a BCA assay and the samples were run on a western blot for the protein of interest as described above.

### Immunocytochemistry (ICC)

SHSY5Y cells were grown in 96-well CellCarrier Ultra plates (Perkin Elmer, 74004), coated with laminin (Sigma, L2020). Cells were then washed in PBS, fixed in 4% PFA (Alfa Aesar, 43368) in PBS for 15mins, washed with PBS and blocked and permeablised for 1 hour in PBS containing 10% donkey serum (DS, Sigma, S30-M) and 0.1% Triton-X-100 (Thermo, T/3751/08) in PBS. Cells were incubated overnight with primary antibodies (Table S2) in PBS containing 0.02% Triton X-100 and 2% DS at 4°C. After washing, secondary antibodies (Table S2) were applied for 1 hour in PBS containing 0.02% Triton X-100 and 2% DS. Cells were stained with Hoescht 33342 (1:2000, Abcam, ab228551) for 10 minutes before the final wash steps. Plates were imaged on the INCell Analyzer 6500 HS with a 40X objective that captured 8 widefield, single plane fields of view per well, and quantitative analysis was performed using Columbus (PerkinElmer).

### Caspase assays

SHSY5Y cells were plated on black clear-bottomed plates (Fisher, 10601442). After 24 hours, cells were stained with NucRed™ Live 647 ReadyProbes™ Reagent (2 drops/ml, Sigma, R37106) and CellEvent™ Caspase-3/7 green ReadyProbes™ Reagent (2 drops/ml, Sigma, R37111) for 1 hour. Staining was removed and cells were then treated with 0.1% DMSO (Sigma, D2650). The plate was imaged and analysed using the IncuCyte.

### mtKeima mitophagy assays

SHSY5Y cells were transduced with mtKeima virus as described above and left for at least a week before analysis. Cells were plated at 10,000 per well in PDL coated 96-well CellCarrier Ultra plates (Perkin Elmer, 16230551), left to settle for 24 hours and then treated with 1uM CCCP for 24 hours in 1% FBS containing media. Cell nuclei were stained with 1:1000 Hoescht 3342 for 10 minutes prior to imaging. Cells were imaged in 1% serum media (Gibco, 31966021) using the INCell Analyzer with a 40X water immersion objective. Randomization of imaging fields was performed through an automated function of the INCell software. In the 488nm channel, which excites mtKeima in neutral environments, mtKeima shows a tubular network-like mitochondrial staining pattern which was labelled as ‘healthy mitochondria.’ In the 561nm channel, which excites mtKeima in acidic environments, the staining was more punctate and was labelled as ‘mitolysosomes.’ Analysis was performed in the Columbus Image Data Storage and Analysis System (Perkin Elmer). Ratiometric analysis of red:green pixel intensity was performed to gain a more robust measure of mitolysosome punctae. The total area of the healthy mitochondria and mitolysosome punctae was quantified and a mitophagy index score calculated by using the ratio of mitolysosome: healthy mitochondrial area.

### qPCR

RNA was isolated using the RNeasy Micro Kit (Qiagen, 74004). cDNA was then reverse transcribed from 200ng RNA using Superscript IV VILO MasterMix (Thermo, 11755250) on the ProFlexTM PCR System (Life Technologies). Triplicate samples containing the appropriate Taqman primers (Ambra1 [Mm00554370_m1] and Beclin1 [Mm01265461_m1]), Applied biosystems Taqman fast advanced mastermix (Thermo, 4444557) and 2 ul cDNA were analysed by qPCR using β-actin (Taqman, [Mm02619580_g1]) as a housekeeping gene and the following programme: 50°C 2 mins, 95°C 20secs, 40 cycles (95°C 1 sec, 60°C 20 secs). QuantStudio7 was used to perform the qPCR and the 2^-DDCt^ method was used for analysis.

### Ki67 proliferation assay

mNSCs were plated onto PEI and laminin coated 96-well CellCarrier Ultra plates (Perkin Elmer, 74004). After 24 hours, a media change was carried out to induce quiescence. Cells were transduced with lentivirus after a following 72 hours. 48 hours post-transduction, cells were fixed with 4% PFA for 15 mins. The fixed mNSCs were washed twice with PBS before permeablisation with 0.1% saponin (Thermo, 47036-50G-F) in 2% BSA (Sigma, A7030) for 1 hour at room temperature. Ki-67 primary antibody diluted 1:200 in 0.1% saponin solution was added overnight at 4°C. Wells were washed in PBS and 100µl of secondary antibody (1:1000, Table S2) diluted in 0.1% saponin solution was added to each well for 1 hour at room temperature. Wells were washed before adding Hoescht 33342 (1:2000) for 10 mins at room temperature. Finally, cells were washed with PBS and incubated at 4°C for imaging on the InCell Analyzer, and quantitative analysis was performed using Columbus.

## Results

### AMBRA1^TAT^ expression protects cells against apoptosis and induces autophagy and mitophagy

AMBRA1^TAT^ is a mutant form of AMBRA1 that cannot interact with DLC1 meaning that AMBRA1 can translocate to the ER and initiate autophagosome formation. Firstly, we confirmed the lentiviral overexpression of AMBRA1^TAT^ in the SHSY5Ys by western blotting. This showed increased levels of AMBRA1 expression in the AMBRA1^TAT^ mutant compared to the control (expressing an empty lentiviral vector, containing a multiple cloning site [MCS] only) **(Figure 1A)**. In addition, we showed that the levels of BECLIN1 phosphorylation were increased in this mutant **(Figure 1B)**. This was expected because AMBRA1^TAT^ should have increased basal levels of autophagy (and therefore BECLIN1 phosphorylation). Next, we carried out Co-IP to assess the interaction between AMBRA1 and DLC1. As expected, the levels of AMBRA1/DLC1 binding were decreased in the AMBRA1^TAT^ mutant cells compared to AMBRA1^WT^ overexpressing cells **(Figure 1C)**.

**Figure 1.**
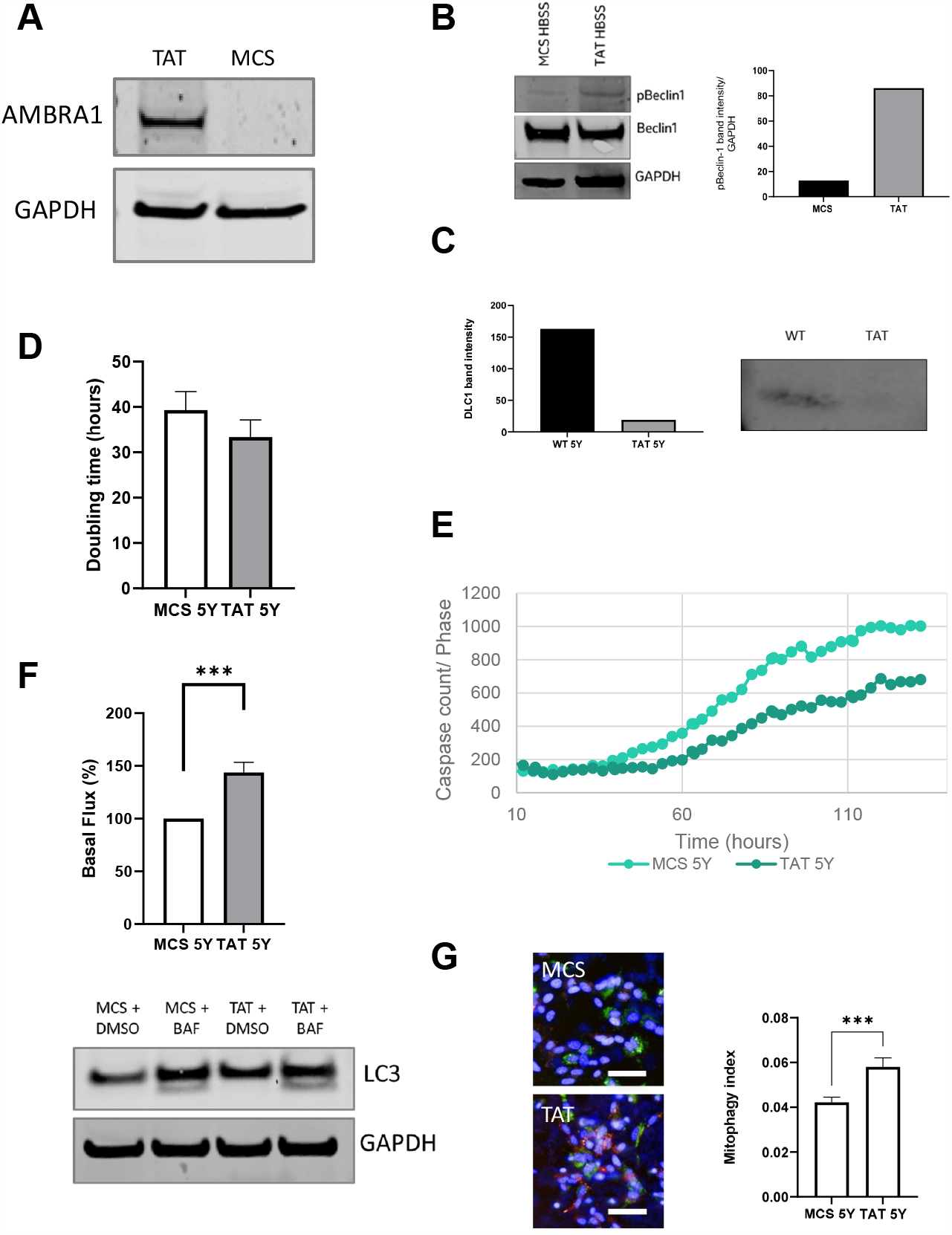
AMBRA1^TAT^ SHSY5Y validation and characterisation. (A) Western blot of AMBRA1^TAT^ and MCS (control) SH-SY5Y cells to show AMBRA1 expression and (B) Western blot and quantification of pBeclin levels. (C) Western Blot and quantification of DLC1 protein expression in Co-IP samples of AMBRA1^TAT^ and MCS SH-SY5Y cells. (D) AMBRA1^TAT^ and MCS SH-SY5Y doubling time measured in the Incucyte over a period of multiple days. (E) AMBRA1^TAT^ and MCS SH-SY5Ys stained with Caspase-3/7 Green and NucRed and imaged in the Incucyte for 130 hours in growth media. Caspase levels were normalised to the phase area of each image. This is a representative graph of three different experimental repeats. (F) Western blot analysis of LC3-II expression in AMBRA1^TAT^ and MCS SH-SY5Y cells treated with growth media and bafilomycin for 4 hours. The data presented is from three separate experiments. Error bars ± SEM.***p<0.0005 (two-tailed unpaired t-test) (G) Normalized quantification of the mitophagy index of AMBRA1^TAT^ and MCS SH-SY5Y cells transduced with mtKeima virus and treated with 1uM CCCP. Scale bar = 50µm. Error bars ± SD.***p<0.0005 (two-tailed unpaired t-test). This is a representative graph of three different experimental repeats.

When the validation of the cell model was complete, we then moved on to characterise the AMBRA1^TAT^ SHSY5Y cells. We saw no change in proliferation between the AMBRA1^TAT^ cells and the MCS control cells **(Figure 1D)**, however expression of the TAT mutant construct was able to protect against apoptosis **(Figure 1E)** in addition to significantly increasing the levels of basal autophagy in the cells **(Figure 1F)** and the mitophagy index, which is indicative of the levels of mitophagy in the cells **(Figure 1G)**. Taken together, these results suggest that blocking the AMBRA1/DLC1 interaction may induce a phenotype that would be beneficial in a neurodegeneration context.

### AMBRA1^WD40^ expression decreases autophagy

AMBRA1^WD40^ cannot interact with DDB1, inhibiting AMBRA1 ubiquitination and its subsequent degradation. There should therefore be more unbound AMBRA1 in the cell to form autophagy and mitophagy related complexes. Western blotting confirmed that AMBRA1^WD40^ SHSY5Y cells had increased levels of AMBRA1 expression compared to the control cells **(Figure 2A)**. In addition, we confirmed that AMBRA1^WD40^ overexpressing cells exhibited lower levels of interaction with DDB1 compared to AMBRA1^WT^ using CoIP **(Figure 2B)**. The AMBRA1^WD40^ overexpressing cells did not show altered proliferation, apoptosis or mitophagy **(Figure 2C-D,F)**. However, AMBRA1^WD40^ overexpression did significantly decrease the levels of basal autophagy in the cells **(Figure 2E)**. Together, these results suggest that targeting the WD40 domain does not produce a phenotype that would be beneficial in a neurodegeneration context.

**Figure 2.**
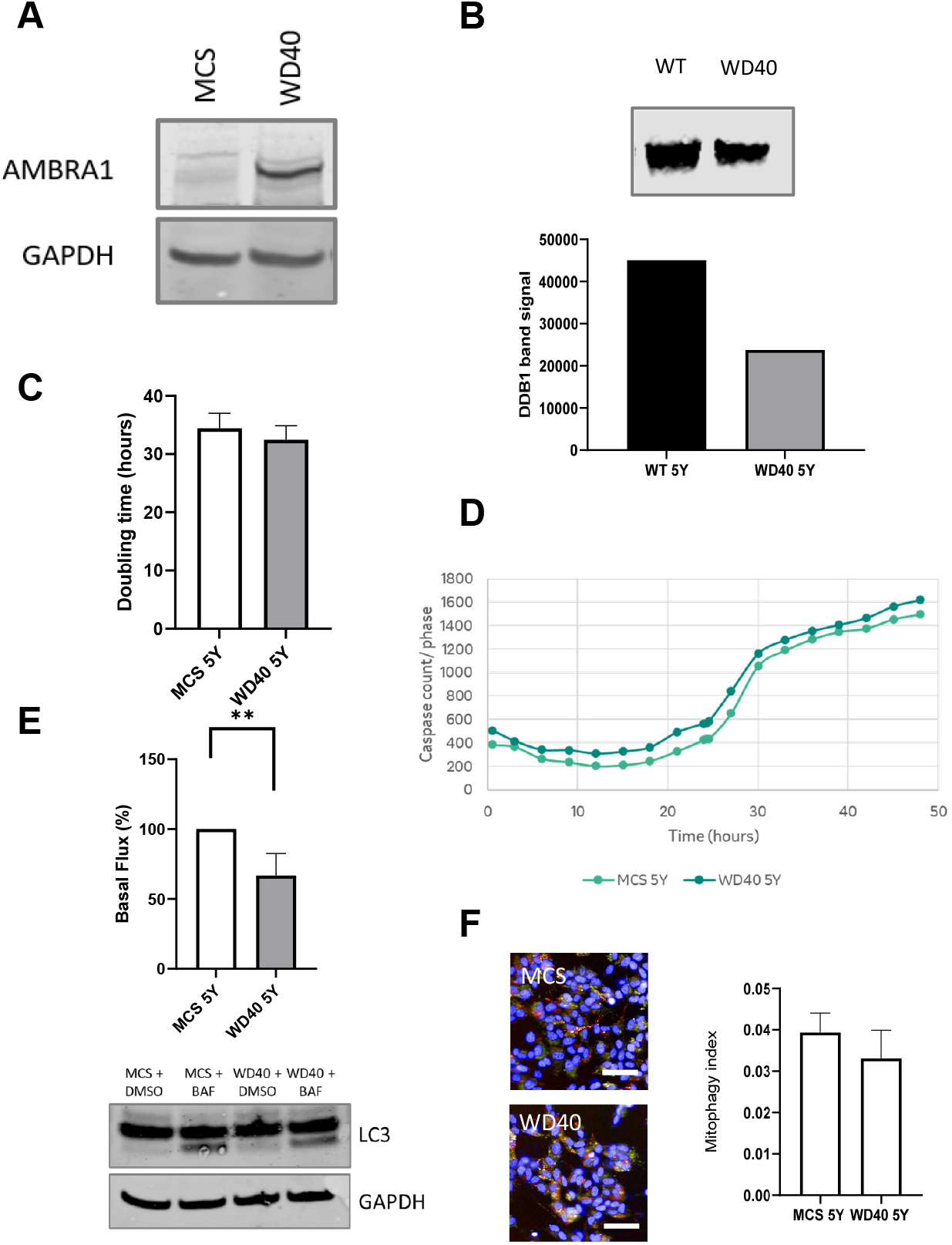
AMBRA1^WD40^ SHSY5Y validation and characterisation. (A) Western blot of AMBRA1^WD40^ and MCS SH-SY5Y cells to show AMBRA1 expression. (B) Western Blot and quantification of DDB1 protein in Co-IP samples of AMBRA1^WD40^ and MCS SH-SY5Y cells. (C) AMBRA1^WD40^ and MCS SH-SY5Y cells doubling time measured in the Incucyte over a period of multiple days. (D) AMBRA1^WD40^ and MCS SH-SY5Y cells stained with Caspase-3/7 Green and NucRed and imaged in the Incucyte for approximately 50 hours in growth media. Caspase levels were normalised to the phase area of each image. (E) Western blot analysis of LC3-II expression in AMBRA1^WD40^ and MCS SH-SY5Y cells treated with growth media and bafilomycin for 4 hours. The data presented is from three separate experiments. Error bars ± SEM.**p<0.005 (two-tailed unpaired t-test). (F) Normalized quantification of the mitophagy index of AMBRA1^WD40^ and MCS SH-SY5Y cells transduced with mtKeima virus and treated with 1uM CCCP. Error bars ± SD.

### AMBRA^S1014^ expression increases autophagy

IKKα has been shown to phosphorylate the S1014 residue of AMBRA1, thus stabilising its interaction with LC3 (Di Rita et al., 2018), therefore we generated a S1014 phospho-mimetic mutant, AMBRA1^S1014^. As expected, there was increased levels of AMBRA1 expression in the AMBRA1^S1014^ mutant cells compared to the MCS cells **(Figure 3A)**. To determine whether the AMBRA1/LC3 interaction was increased, we carried out CoIP which confirmed that the AMBRA1^S1014^ overexpressing cells had increased levels of interaction between LC3 and AMBRA1 compared to AMBRA1^WT^ overexpressing cells **(Figure 3B)**. Expression of AMBRA1^S1014^ did not affect proliferation, apoptosis or mitophagy **(Figure 3C-D,F)** but did increase the levels of basal autophagy **(Figure 4E)**. Therefore, the only mutant that was able to affect apoptosis, autophagy and mitophagy in beneficial ways was the TAT mutant.

**Figure 3.**
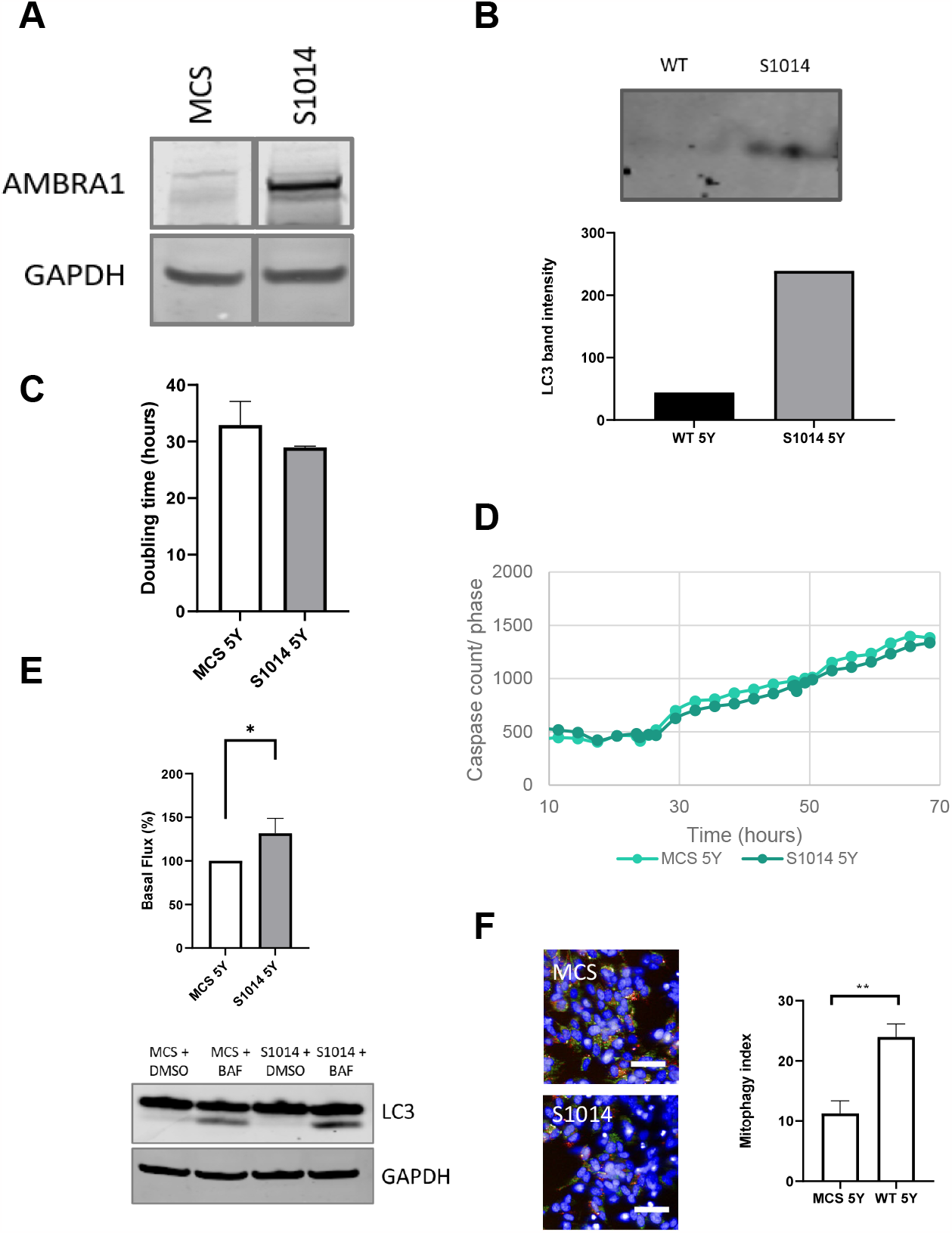
AMBRA1^S1014^ SHSY5Y validation and characterisation. (A) Western blot of AMBRA1^S1014^ and MCS SH-SY5Y cells to show AMBRA1 expression. (B) Western Blot and quantification of LC3 protein in Co-IP samples of AMBRA1^S1014^ and MCS SH-SY5Y cells. (C) AMBRA1^S1014^ and MCS SH-SY5Y cells doubling time measured in the Incucyte over a period of multiple days. (D) AMBRA1^S1014^ and MCS SH-SY5Y cells stained with Caspase-3/7 Green and NucRed and imaged in the Incucyte for approximately 70 hours in growth media. Caspase levels were normalised to the phase area of each image. (E) Western blot analysis of LC3-II expression in AMBRA1^S1014^ and MCS SH-SY5Y cells treated with growth media and bafilomycin for 4 hours. The data presented is from three separate experiments. . Error bars ± SEM. *p<0.05 (two-tailed unpaired t-test). (F) Normalized quantification of the mitophagy index of AMBRA1^S1014^ and MCS SH-SY5Y cells transduced with mtKeima virus. Error bars ± SD. Two two-tailed unpaired t-tests where *p<0.05.

**Figure 4.**
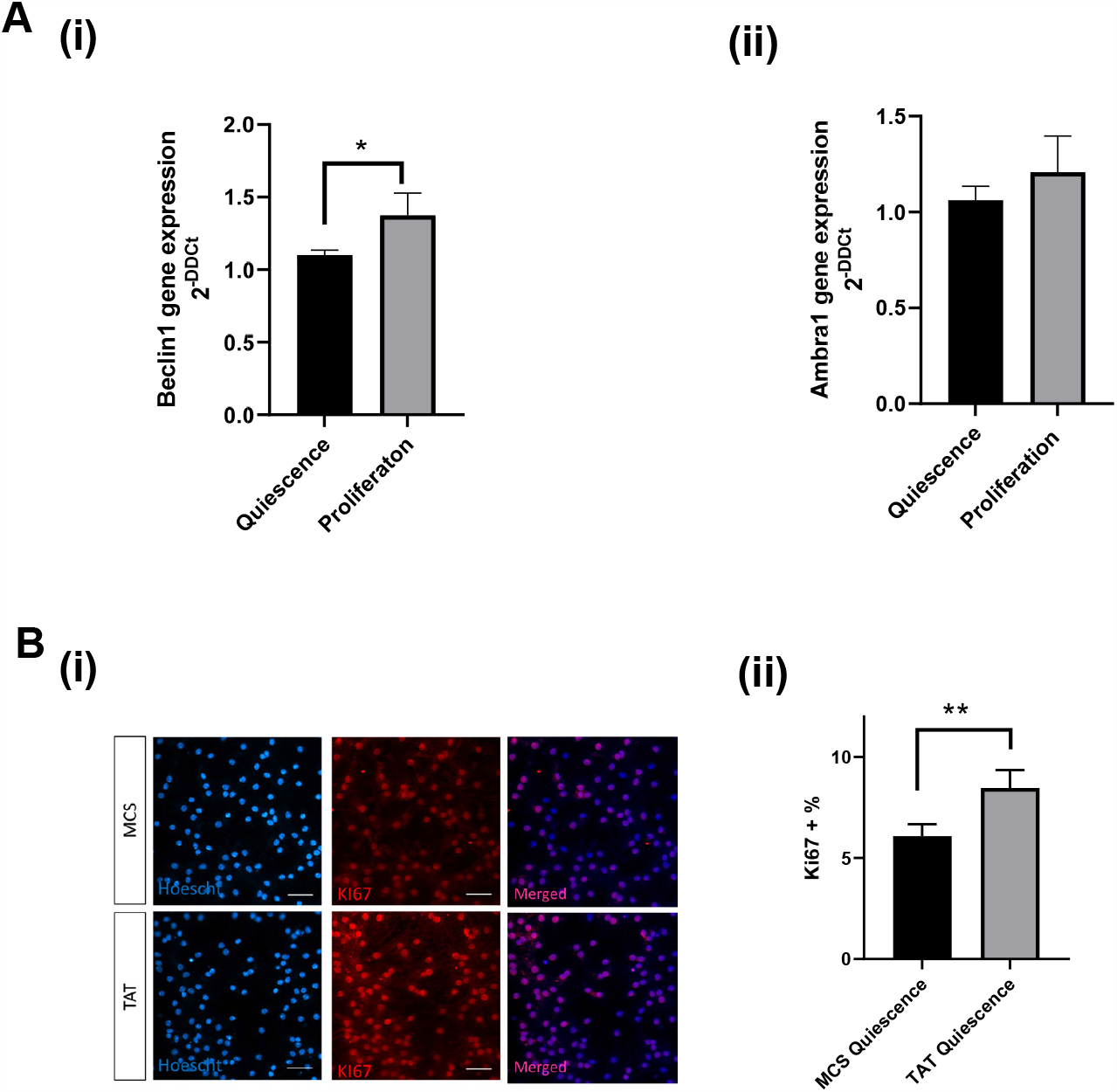
AMBRA1^TAT^ overexpression in mouse neural stem cells. (A) Quantification of the fold change of Beclin1 (i) and Ambra1 (ii) transcript expression in mNSCs which have been reactivated from quiescence to proliferation. (B)(i)-(ii) Immunostaining of Hoescht (blue) and Ki-67 (red) in proliferating and quiescent mNSCs. Quiescence was induced using BMP-4. Scale bar = 50µm. Images taken at 40x magnification. Averages taken from 8 fields of view. Error bars ± SEM. Two-tailed unpaired t-test where *p<0.05 and **p<0.01.

### AMBRA1^TAT^ expression enhances the reactivation of quiescent mouse neural stem cells

AMBRA1 has been implicated in neurodevelopment and recent studies have also implicated AMBRA1-mediated autophagy as a key process during adult neurogenesis (Gillotin et al., 2021). Indeed, regulation of autophagy has emerged as a key signalling pathway to modulate the balance between quiescent and proliferative mNSCs and dysregulation of this mechanism is thought to be one underlying causes of cognitive impairment in several neurodegenerative diseases (Gillotin et al., 2021). We therefore investigated whether AMBRA1 affects the shift from quiescence to proliferation in a well-established *in vitro* model (Martynoga et al., 2013). First, we assessed the levels of *Ambra1* and *Beclin1* expression in mNSCs undergoing the switch from quiescence to proliferation. *Beclin1* gene expression significantly increased during this transition **(Figure 4A(i))** and *AMBRA1* showed a similar trend, although not reaching statistical significance **(Figure 4A(ii))**. Since lysosome activation was shown to promote reactivation of quiescent NSC *in vitro and vivo*, (Leeman et al., 2018) we next used the AMBRA1^TAT^ mutant, due to its gain-of-function related to autophagy, to assess whether activation of AMBRA1 could enhance the cellular transition from quiescence to proliferation. Interestingly, overexpression of AMBRA1^TAT^ in mNSCs significantly increased the proportion of Ki-67^+^ proliferative cells compared to the MCS control **(Figure 4B(i-ii))**. These results suggest that activation of AMBRA1 by disruption of the AMBRA1/DLC1 interaction may help promote the reactivation of quiescent NSCs, a major roadblock to sustain adult neurogenesis in neurodegenerative diseases.

### AMBRA1^ACTA^ expression induces mitophagy and protects cells against apoptosis

AMBRA1^ACTA^ is a form of AMBRA1 with a mitochondrial localisation sequence which should induce translocation of AMBRA1 to the outer mitochondrial membrane (OMM) to induce mitophagy. In order to validate that AMBRA1^ACTA^ translocates to the mitochondria we performed immunofluorescent staining of AMBRA1 and TOM20, an OMM marker, which demonstrated that these two proteins were both localised to the mitochondria in the AMBRA1^ACTA^ but not the control cells **(Figure S1A)**. AMBRA1^ACTA^ overexpressing cells also exhibited increased AMBRA1 protein expression **(Figure S1B)** in addition to an unchanged proliferation rate **(Figure S1C)**, protection against apoptosis **(Figure S1D)**, increased levels of autophagy **(Figure S1E)** and significantly increased mitophagy **(Figure S1F)**. In conclusion, this mutant has desirable effects on apoptosis and autophagy/mitophagy.

## Discussion

AMBRA1 is a scaffold protein that performs as a platform for various different molecular processes, in particular it acts at the crossroads between autophagy and apoptosis (Fimia et al., 2013). Furthermore, processes that are influenced by AMBRA1 have been implicated in neurodevelopment and neurogenesis (Vazquez et al., 2012, Yazdankhah et al., 2014, Wang et al., 2022), making AMBRA1 activation a potential therapeutic strategy for neurodegenerative disease. However, a chemical strategy for AMBRA1 activation is yet to be established.

Di Rita et al., (2018b) previously showed that AMBRA1 localisation to the OMM via the overexpression of AMBRA1^ACTA^ in HeLa cells can stimulate mitophagy. Our findings corroborated this, demonstrating significantly increased levels of both autophagy and mitophagy in SHSY5Y cells overexpressing this construct. Interestingly, overexpression of this mutant construct also protected the cells against apoptosis, which could be due to the increased mitophagy decreasing the levels of damaged mitochondria and thus improving the overall health of the cells.

The AMBRA1^WD40^ mutant construct was designed to prevent the interaction between AMBRA1 and DDB1. Since the DDB1/Cullin4 complex ubiquitinates AMBRA1 this mutant should have lower levels of AMBRA1 ubiquitination and subsequent degradation, similar to what was observed by Antonioli et al. (2014). We hypothesised that the AMBRA1 accumulation that results would allow the formation of complexes involved in autophagy and mitophagy and thus stimulate these processes. However, autophagic flux was found to be lower in the AMBRA1^WD40^ mutant cells and there was no difference in any of the other parameters compared to the control. These findings were unexpected and suggest that alternative complexes, such as the RNF2/WASH complex, that initiates proteasomal degradation (Xia et al., 2014) of AMBRA1, do this to a greater extent in the absence of the interaction between AMBRA1 and DDB1/CULLIN4. The WD40 domain of AMBRA1 is the only domain with a known structure and would therefore be the most straightforward to target pharmacologically. Despite this, our results would not support this as a viable drug strategy for neurodegeneration since AMBRA1/DDB1 protein-protein interaction inhibitor (PPI) compounds would likely block autophagy rather than promoting it.

Overexpression of the AMBRA1^S1014^ mutant construct in SHSY5Y cells affected autophagy but none of the other parameters tested. These findings were expected in that increasing the interaction between AMBRA1 and LC3 should increase autophagy, but you might also expect this to increase the levels of mitophagy since the AMBRA1/LC3 interaction is also involved in this process. Indeed, Di Rita et al., (2018b) observed increased mitophagy in this phospho-mimetic mutant. However, this was in HeLa cells which do not have an active PARKIN-dependent mitophagy pathway, and therefore PARKIN-independent mitophagy such as that coordinated by AMBRA1/HUWE1 is likely to play a more predominant role. We used SHSY5Y cells in preference to HeLa cells due to the differences in the levels of AMBRA1 expression between the two cell lines. HeLa cells have high levels of AMBRA1 protein expression compared to SHSY5Y cells (data not shown) in which the levels are extremely low and therefore endogenous AMBRA1 is not likely to interfere with the mutant AMBRA1 constructs that we overexpressed as it would in HeLa cells.

Of all the mutant constructs characterised, AMBRA1^TAT^ showed the most promise in that its overexpression significantly increased both autophagy and mitophagy whilst simultaneously protecting cells from apoptosis. In addition, there was no effect on cellular proliferation which ensures the tumorigenesis risk of this activation strategy is low. AMBRA1^ACTA^ mutants exhibited the same phenotype however this does not represent a viable pharmacological strategy. It has previously been shown that AMBRA1^TAT^, which exhibits reduced binding to DLC1, increases the levels of autophagy in cells (Di Bartolomeo et al., 2010). However, we show for the first time that this mutant can also induce mitophagy. This is unsurprising since the molecular mechanisms are highly conserved between the two processes.

Moreover, we show here that mNSCs transduced with the AMBRA1^TAT^ mutant construct exhibit enhanced reactivation from quiescence to proliferation. This switch is central to adult neurogenesis as most NSCs remain in quiescence and their inability to switch back to a proliferative state increases with age and in several neurodegenerative diseases (Gillot et al., 2021). In this context, AMBRA1^TAT^ is likely increasing autophagy, similar to what we observed in the SHSY5Y cells, which could supply degradable substrates to the lysosomes to activate quiescent NSCs reinforcing a well-established as process to enhance adult neurogenesis.

Since th AMBRA1^TAT^ mutant represents a potential PPI strategy between AMBRA1/DLC1, we suggest that this approach in particular should be pursued if AMBRA1 activation is to be used as a drug discovery strategy. The limitation to this approach currently is the lack of a known structure for the intrinsically disordered regions of AMBRA1. Therefore, future work should focus on determining the AMBRA1 structure, in particular the interaction region between AMBRA1 and DLC1, and designing PPI compounds to disrupt this interaction and ultimately, to interfere with molecular mechanisms promoting neurodegeneration thus providing therapeutic benefit.

## Acknowledgements

The project was funded solely by Merck & Co., Inc., Rahway, NJ, USA.

## Declaration of Interests

All authors are employees and shareholders of Merck & Co., Inc., Rahway, NJ, USA.

## Tables

**Table S1.**
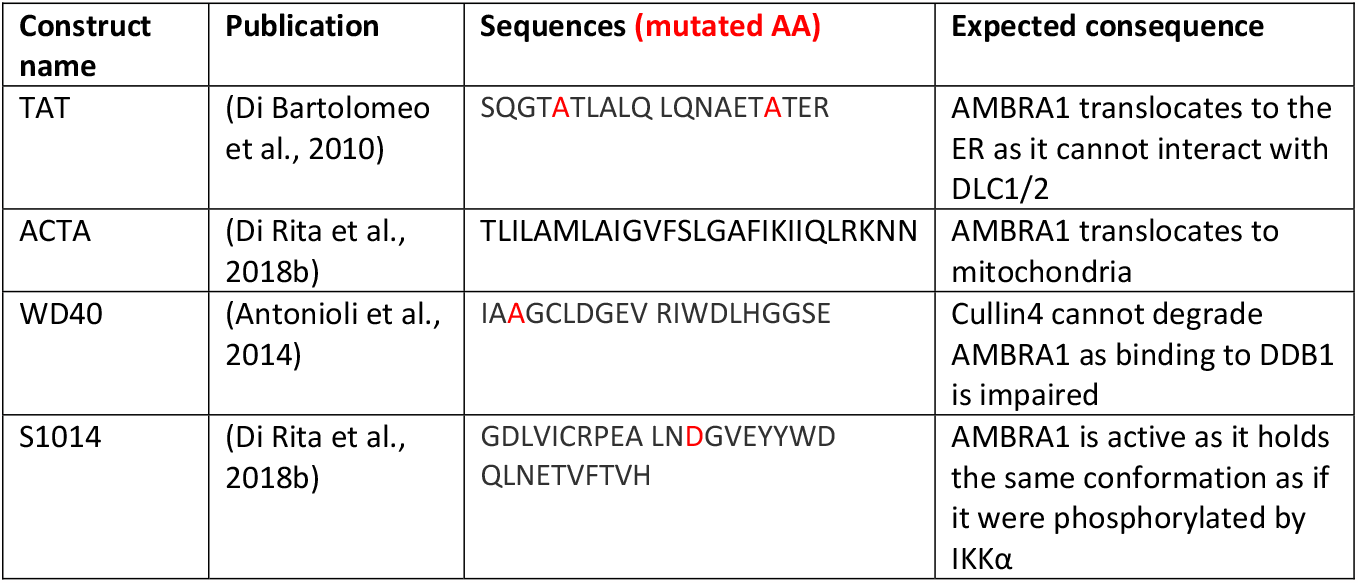
AMBRA1 mutants.

**Table S2.** Antibodies.

| Antibody | Company, catalog number | Dilution |
| --- | --- | --- |
| Anti-GAPDH Mouse Monoclonal Antibody | Abcam, ab8245 | 1:4,000 |
| Recombinant Anti-DYNLL1/PIN Rabbit Monoclonal antibody | Abcam, ab51603 | 1:200 |
| Recombinant Anti-DDB1 Rabbit Monoclonal Antibody | Abcam, ab109027 | 1:200 |
| Recombinant Anti-LC3B Antibody | Abcam, ab192890 | 1:500 |
| pBeclin-1 Rabbit Monoclonal Antibody | CST, 84966S | 1:500 |
| Anti-LC3B Rabbit Polyclonal Antibody | Novus Bio, NB100-2220 | 1:500 |
| AMBRA1 Rabbit Polyclonal Antibody | Proteintech, 13762-1-AP | 1:100 |
| Total Beclin Rabbit PolyAB Antibody | Proteintech, 11306-1-AP | 1:500 |
| Ki-67 Rat Monoclonal Antibody | ThermoFisher, 14-5698-82 | 1:200 |
| Donkey anti-rabbit mouse | Fisher, A21206 | 1:1,000 |
| Donkey anti-mouse red | Fisher, A31571 | 1:1,000 |
| IRDye<sup>®</sup> 800CW Donkey anti-Rabbit IgG | Licor, 15590485 | 1:4,000 |
| IRDye<sup>®</sup> 680 RD Donkey anti-mouse IgG | Licor, 15550535 | 1:4,000 |

---

**Figure S1.**
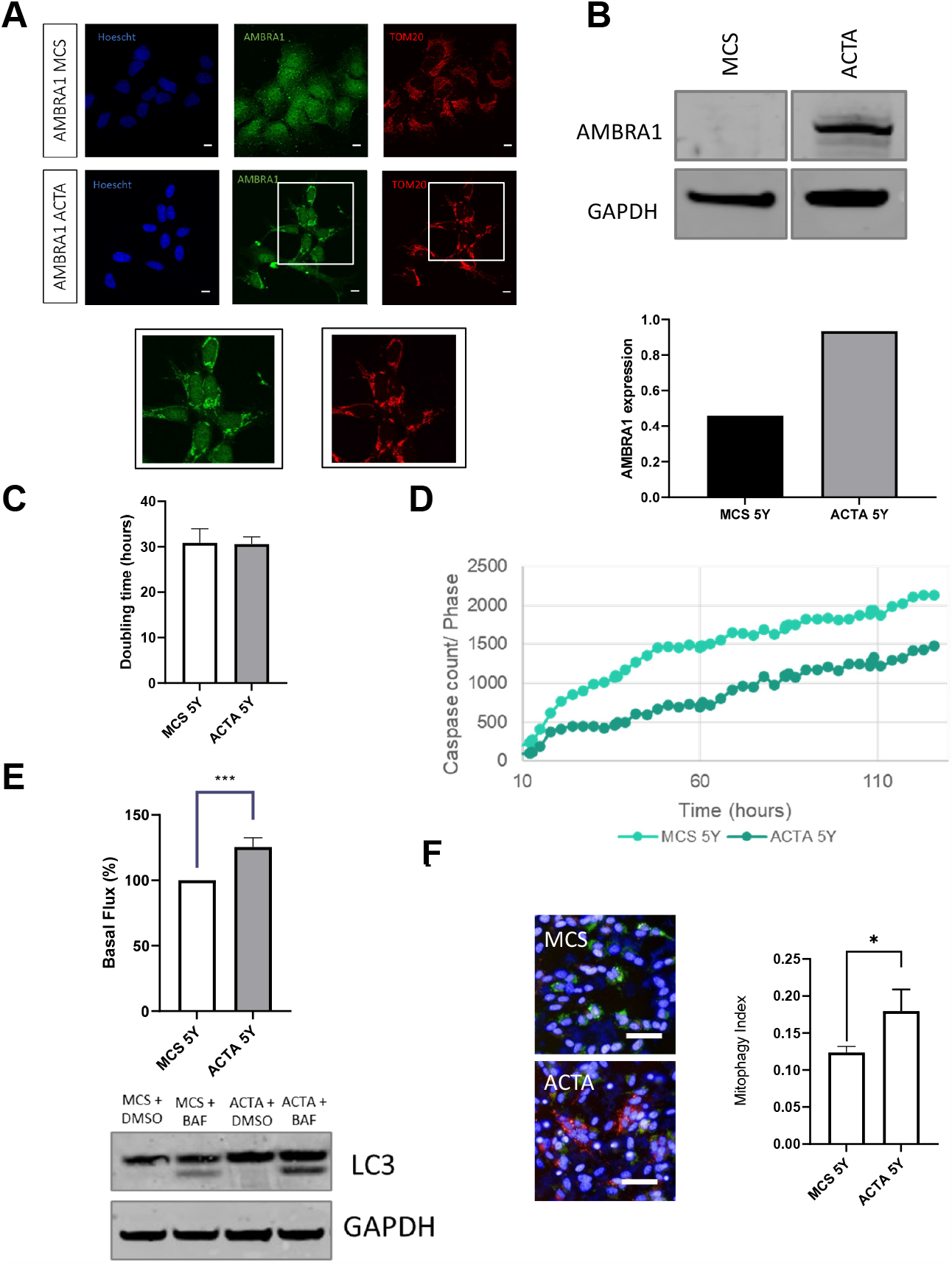
AMBRA1^ACTA^ SHSY5Y validation and characterisation. (A) Immunostaining of Hoescht (blue), TOMM20 (green) and AMBRA1 (red) in AMBRA1^ACTA^ and MCS SH-SY5Y cells. Scale bars =10um (B) Western blot of AMBRA1^ACTA^ and MCS SH-SY5Y cells to show AMBRA1 protein expression. (C) AMBRA1^ACTA^ and MCS SH-SY5Y doubling time measured in the Incucyte over a period of multiple days. (D) AMBRA1^ACTA^ and MCS SH-SY5Y cells stained with Caspase-3/7 Green and NucRed and imaged in the Incucyte for approximately 130 hours in growth media. Caspase levels were normalised to the phase area of each image. (E) A western blot analysis of LC3-II expression in AMBRA1^ACTA^ and MCS SH-SY5Y cells treated with growth media and bafilomycin for 4 hours. Error bars ± SEM. (F) Normalized quantification of the mitophagy index of AMBRA1^ACTA^ and MCS SH-SY5Y cells transduced with mtKeima virus and treated with 1uM CCCP. Scale bar = 50µm. Error bars ± SD. Two two-tailed unpaired t-tests where *p<0.05, .***p<0.0005 . This is a representative graph of three different experimental repeats.

